# Anti-tumor effects of a novel cell penetrating peptide-based therapeutic approach to target Lactate Dehydrogenase C (LDHC) in triple negative breast cancer

**DOI:** 10.1101/2025.03.11.641612

**Authors:** Hanan Qasem, Adviti Naik, Tricia Gomez, Janarthanan Ponraj, Umar Jafar, Martin Sikhondze, Remy Thomas, Khaled A Mahmoud, Julie Decock

**Affiliations:** College of Health and Life Sciences (CHLS), Hamad Bin Khalifa University (HBKU), Qatar Foundation (QF), Qatar; Translational Oncology Research Center, Qatar Biomedical Research Institute (QBRI), Hamad Bin Khalifa University (HBKU), Qatar Foundation (QF), Qatar; Qatar Environment and Energy Research Institute (QEERI), Hamad Bin Khalifa University (HBKU), Qatar Foundation, Qatar

**Keywords:** LDHC, Lactate Dehydrogenase C, CPP, cell penetrating peptide, triple negative breast cancer, xenograft zebrafish

## Abstract

**Background:** Lactate Dehydrogenase C (LDHC) is a promising candidate for therapeutic targeting thanks to its highly tumor-specific expression, immunogenicity, and pro-tumorigenic functions. Aberrant LDHC expression is associated with poor clinical outcomes in multiple cancers, including breast cancer. However, no specific LDHC inhibitors are currently available, highlighting the need for novel strategies to selectively target LDHC in tumor cells. This study explores the anti-tumor potential of cell-penetrating peptides (CPPs) to target LDHC in triple negative breast cancer (TNBC).

**Methods:** Four CPPs were evaluated for their ability to deliver LDHC siRNA to tumor cells, including the positively charged 10R peptide (10R) and three bifunctional peptides containing the integrin αvβ3 recognition motif Arg-Gly-Asp (RGD): 10R-RGD, cyclicRGD-10R (cRGD-10R), and internalizing RGD-10R (iRGD-10R). We characterized the physicochemical properties of all CPP:siRNA complexes, and determined their serum stability, cytotoxicity, cellular uptake, and *LDHC* silencing efficiency in vitro. The anti-tumor effects and cytotoxicity of cRGD-10R:siRNA and iRGD-10R:siRNA complexes were further assessed in a TNBC xenograft zebrafish model.

**Results:** All four CPPs formed stable nanocomplexes with favorable safety profiles. The 10R-RGD and cRGD-10R peptides demonstrated the most efficient LDHC knockdown, reduced the clonogenic ability of TNBC cells and enhanced their treatment response to the chemotherapeutic drug olaparib in vitro. Treatment of TNBC xenograft zebrafish with 10R-RGD:siRNA and cRGD-10R:siRNA complexes significantly reduced tumor burden without inducing major toxicity. **Conclusion** Our findings demonstrate that CPP-based siRNA delivery provides a novel and safe approach to target LDHC, either as a monotherapy or in combination with common anti-cancer drugs, to enhance treatment outcomes.

## INTRODUCTION

Breast cancer is the most prevalent cancer and leading cause of mortality among women worldwide [1]. Triple negative breast cancer (TNBC), characterized by the lack of expression of the estrogen receptor, progesterone receptor, and Human Epidermal Growth Factor Receptor 2 (Her2), represents 10-20% of breast cancer cases, primarily affects young women and is associated with a poor prognosis and high metastatic potential [2,3]. Currently, the treatment options for patients with TNBC include chemotherapy, radiotherapy and more recently immunotherapy. While TNBC patients exhibit early pathological responses to chemotherapy, unfortunately, a significant portion of patients develop resistance and disease recurrence.

In recent years, cancer treatment is shifting away from a one-size-fits-all approach towards precision medicine, where traditional treatments such as surgery, chemotherapy, and radiotherapy are complemented by personalized drug therapies [2,4]. Recent studies suggest that combining molecular targeting with chemotherapy represents a promising approach to treat TNBC [5]. For instance, the addition of mTOR inhibitors, such as temsirolimus or everolimus to doxorubicin and bevacizumab has significantly improved the objective response rate of TNBC patients with tumors that display aberrant activation of the PI3K/mTOR pathway [6]. In addition, combination treatment of carboplatin with the PARP inhibitor olaparib is associated with an overall response rate (ORR) of 88% in TNBC patients carrying BRCA mutations. In this context, specific targeting of cancer/testis antigens (CTAs) has gained interest thanks to their highly tumor-specific expression, immunogenic properties and multifaceted roles in promoting cancer hallmarks [7–10]. Aberrant expression of Lactate Dehydrogenase C (LDHC) is observed in different types of cancer, where it is associated with tumor progression, metastasis, and poor prognosis [11–13]. Previously, we demonstrated that knockdown of LDHC reduces long term survival of breast tumor cells, in particular of TNBC cells, through dysregulation of cell cycle progression and impairment of the DNA damage response pathway [14]. LDHC can thus be included in the expanding group of CTAs involved in regulating genomic integrity. Moreover, we found that targeting LDHC greatly improved treatment response to commonly used DNA damage response-related drugs such as cisplatin and olaparib.

Gene therapy using RNA interference (RNAi)-based drugs has shown remarkable progress with numerous tumor-related RNAi drugs undergoing clinical trials [15]. For instance, RNAi therapeutics targeting anti-apoptosis genes, oncogenes and tumor signaling molecules such as Bcl-2, MYC, KRAS, AKT1 and STAT3 have entered phase II trials. Despite promising preclinical outcomes, RNAi-based therapy has yet to transition into clinical practice as several challenges remain to be addressed including stability, targeting ability, off-target effects, and toxicity. Advances in the development of cell penetrating peptides (CPPs) as delivery systems for RNAi-based therapy have aided to address some of these challenges. The use of CPPs improves RNAi serum stability and internalization efficiency. Additionally, CPPs containing the arginine-glycine-aspartic acid (RGD) tripeptide enhance tumor specificity through binding of integrins αvβ3 (INTαVβ3) [16]. The use of CPPs has demonstrated efficient delivery of siRNA cargo with notable anti-tumor activity. For example, cyclic-RGD:siEGFR reduced EGFR tumor expression by 50% and significantly decreased tumor size in glioblastoma (U87MG) xenograft mice [17]. Furthermore, multiple studies have reported that CPPs incorporating an internalizing RGD peptide (iRGD) enhance tumor penetration and improve anti-cancer therapeutic effects [18,19]. In this study, we investigated the delivery efficacy, anti-tumor activity and safety of four distinct CPPs targeting LDHC including polyarginine (10R), 10R-linear RGD (10R-RGD), cyclic-RGD-10R (cRGD-10R) and internalizing RGD-10R (iRGD-10R). All CPPs formed uniform, positively charged RNAi-nanocomplexes with good serum stability, efficient cellular uptake and effective LDHC knockdown in triple negative breast cancer cell lines. Each of the CPP:siRNA nanocomplexes was associated with a favorable safety profile in non-cancerous and cancerous cell lines. Administration of particularly the 10R-RGD and cRGD-10R peptide:siLDHC complexes decreased long term tumor cell survival and improved chemotherapy responses in vitro. Moreover, 10R-RGD and cRGD-10R based delivery of siLDHC significantly reduced tumor burden in TNBC xenograft zebrafish in the absence of toxicity.

## METHODS

### Cell penetrating peptides (CPPs) and siRNA

CPPs were synthesized by ThermoFisher scientific (USA) using solid-phase peptide synthesis and purity was determined by high-performance liquid chromatography (HPLC). CPP sequences, purity and molecular weight are listed in **Table 1**. CPPs were resuspended in RNase/DNase free water at 1mg/ml. Accell siLDHC#1 (A-008759-14-0020), siLDHC#2 (A-008759-15-0020) and siCTRL1 (D-001910-20) were obtained from Dharmacon (Lafayette, CO, USA) and resuspended at 100 µM.

**Table 1:**
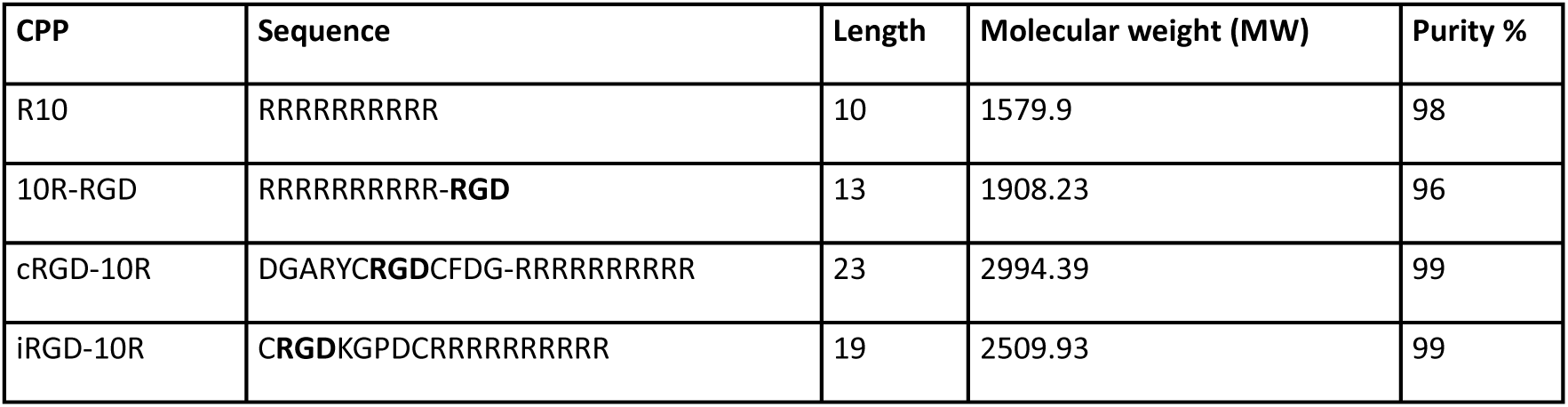
The biochemical properties of CPPs.

### CPP:siRNA complex preparation

CPP (1 mg/ml stock) and siRNA (100 µM stock) were mixed at various peptide:siRNA molar ratios (2.5:1, 5:1, 10:1) in 50 µl of Ultra-Pure DNase/RNase free water. After 45 min incubation at room temperature, Opti-MEM I Reduced Serum medium (Gibco, #11058-021) was added to achieve a final siRNA concentration of 200nM and CPP concentration of 30-60 nM.

### Gel retardation assay

CPP:siRNA complex solutions at various peptide:siRNA molar ratios were supplemented with 1.25x formamide loading dye, and resolved on a 20% native polyacrylamide gel electrophoresis (PAGE) for 120 minutes at 120 V using 1X Tris-Boric-EDTA (TBE) buffer, alongside a 100bp DNA ladder (ThermoFisher Scientific,#SM024). The gels were stained with SYBR gold (Invitrogen, S11494) for 20 mins and analyzed under UV light using ChemiDoc (Bio-Rad). Unconjugated or naked siRNA was used as control.

### Physicochemical characterization of CPP:siRNA nanocomplexes

Particle size, zeta potential, polydispersity index and conductivity were measured for each CPP:siRNA complex, diluted in 1ml Ultra-Pure DNAse/RNAse free water, using a Zetasizer Ultra ZSU5700 (Malvern Instruments, Inc., Worcestershire, UK) at a wavelength of 677nm with a constant angle of 90° at 37⸰C. Additionally, complex morphology and size was analyzed by transmission electron microscopy using the TALOSF200X microscope (ThermoFisher scientific). Briefly, 10-µL of sample suspension was applied onto a 300-mesh carbon-coated copper grid, negatively stained with uranyl acetate (Electron microscopy science, #22405), and air-dried prior to collecting images using the Talos F200C Transmission Electron Microscope (Thermo Fisher Scientific).

### Serum stability assay

CPPs were complexed with siLDHC#2 at 5:1 peptide:siRNA molar ratio for 45 min at room temperature. Next, CPP:siRNA complexes and naked siRNA were incubated at 37 °C with 50% normal human serum (Invitrogen,#31876) or Ultra-Pure DNAse/RNAse free water for 6, 12 or 24 hours and frozen at −20 °C. Samples were centrifuged at 10,000 rpm for 5 min at 4°C and pellets were resuspended in 15 μl of Ultra-Pure DNAse/RNAse free water. Finally, samples were incubated with proteinase K (0.006 mM), CaCl_2_ (0.3 mM) and Tris-HCl (3 mM, pH = 7.0) for 5.5-6 hours at 37 °C. Formamide loading buffer was added to the samples after which electrophoresis was performed using native 20% PAGE for 120 min at a constant voltage of 120 V. Gels were stained with SYBR Gold (Invitrogen, S11494) for 20 min, and analyzed using ChemiDoc^TM^ XR (Bio-Rad).

### Cell culture

MDA-MB-453, MDA-MB-468, BT-549, DU4475, HUVEC and IMR-90 were purchased from the American Tissue Culture Collection (ATCC). MDA-MB-468 and MDA-MB-453 cells were maintained in Dulbecco’s Minimum Essential Media (DMEM, Gibco, #10569-010) supplemented with 10% v/v fetal bovine serum (FBS, Gibco, #10082-147), 50 U/mL Penicillin and 50 µg/mL Streptomycin (Gibco, #15140-122). BT-549 cells were cultured in ATCC-formulated Roswell Park Memorial Institute (RPMI)-1640 medium (Gibco, A10491-01) supplemented with 10% (v/v) FBS (Gibco, #10082-147), 50 U/mL penicillin and 50 µg/mL streptomycin (Gibco, #15140-122), and 0.023 IU/mL insulin (Sigma-Aldrich, #11070-73-8). DU4475 cells were maintained in ATCC-formulated Roswell Park Memorial Institute (RPMI)-1640 medium (Gibco, A10491-01) supplemented with 10% (v/v) FBS (Gibco, #10082-147, 50 U/mL Penicillin and 50 µg/mL Streptomycin (Gibco, #15140-12). HUVEC cells were maintained in EBM-2 (Lonza, cc-3156) supplemented with EGM^TM^-2 singleQuots supplements (Lonza, # CC-4176) and using collagen coated flask (Thermoscientific, #132606). IMR-90 cells were cultured in Minimum Essential Media (MEM, Gibco, #41090-028) supplemented with 10% v/v FBS (Gibco, #10082-14), 50 U/mL Penicillin and 50 µg/mL Streptomycin (Gibco, #15140-122), 1% sodium pyruvate (Gibco, #11360-039), 1% non-essential amino acids (Gibco, #11140-050). All cell lines were maintained at 37 °C and 5% CO_2_ in a humidified incubator. Regular mycoplasma testing was conducted using a polymerase chain reaction (PCR)-based detection assay. Early passage cells (< P10) were used for all experiments.

### Cellular uptake of CPP:siRNA nanocomplexes

To enable visualization of cellular uptake of the CPP:siRNA complexes in cancer cells, siRNA was pre-labeled with Cy™3 using the Silencer™ siRNA Labeling Kit (ThermoFisher scientific, #2960050). A total of 1×10^5^ MDA-MB-468 breast cancer cells were plated per well in a 12-well plate and left overnight at 37 °C and 5% CO2. Next, cells were washed once with DPBS Gibco, #14190-094) and pre-incubated in Opti-MEM I Reduced Serum medium (Gibco, #11058-021) for 30 min after which 500µl of CPP:siRNA (400nM siRNA in Opti-MEM I Reduced Serum medium) was added to each well. After 6 hours, 500ul of complete, antibiotic-free DMEM media was added (final siRNA concentration= 200 nM). In parallel, cells were incubated with either 100pmol of siRNA alone (naked siRNA control) or were transfected with siRNA using lipofectamine 3000 (Thermo Fisher, #3000-001) according to the manufacturer’s guidelines. After 72 hours, cellular uptake of CPP:siRNA-Cy3 complexes was visualized using the Olympus IX73 microscope at 10X magnification.

### Flow cytometry analysis of integrin αvβ3 expression

A range of breast cancer and non-cancerous cell lines with varying expression of integrin αvβ3 was selected to assess RGD-mediated cellular uptake and toxicity of CPP:siRNA complexes. Integrin αvβ3 expression was analyzed using flow cytometry, real time qRT-PCR, and western blotting. For flow cytometry, a total of 1×10^5^ cells was incubated with 100 µl stain buffer (BD Bioscience, #554656), supplemented with 5μl of human FcR Blocking reagent (Miltenyi Biotec, #130-059-901). After 15 min incubation at 4°C, cells were incubated with mouse anti-human integrin αvβ3 BV421 conjugated antibody (BD Biosciences, #744088) at 1:20 for one hour followed by two washes with DPBS (Gibco, #14190-094) at 300 x g for 5 min at room temperature. Finally, integrin αvβ3 expression was analyzed on the BD LSRFortessa X-20 instrument (BD Biosciences). For each sample 10,000 events were recorded, and further analysis was performed using FlowJo™ Software (BD Biosciences, version 10.8).

### Expression analysis of *LDHC* and *integrins* using quantitative real-time reverse transcription polymerase chain reaction (qRT-PCR)

To assess the effect of CPP:siRNA treatment on *LDHC* expression, total RNA was isolated 72 hours after treatment using the RNeasy Mini kit (Qiagen, #74106). In addition, total RNA was extracted from a range of breast cancer and non-cancerous cell lines to determine the expression of *integrin αv* and *integrin β3*. RNA quantity and purity was assessed using A260/A280 and A260/A230 measurements on a Nanodrop2000 spectrophotometer (ThermoFisher Scientific). Next, cDNA was synthesized from 1 µg of total RNA using the M-MLV Reverse Transcriptase kit (Invitrogen, #28025-013) according to the manufacturer’s guidelines. *LDHC* expression was quantified using a specific 5′FAM-3′MGB TaqMan gene expression primer/probe set (Hs00255650_m1, Applied Biosystems, Foster City, CA, USA). The mRNA expression of integrin αv and integrin β3 was quantified using 100ng of cDNA, specific SYBR-based qPCR primers (integrin αv F: 5-GGGACTCCTGCTACCTCTGT-3, integrin αv R: 5-GAAGAAACATCCGGGAAGACG-3, integrin β3 F: 5-ACTGGCAAGGATGCAGTGAA-3 and integrin β3 R: TTGGACACTCTGGCTCTTC-3) and the PowerUp SYBR Green master mix (Applied Biosystems, #A25742). All reactions were performed on the QuantStudio 7 Real-time PCR instrument (Applied Biosystems). Expression levels were normalized to the housekeeping gene *RPLPO* (TaqMan primer/probe 4333761F or SYBR primers F: TCCTCGTGGAAGTGACATCG, R: TGGATGATCTTAAGGAAGTAGTTGG) and differential gene expression was calculated using the 2^−ΔΔCt^ method.

### Western blotting of LDHC and integrins

Western blotting was used to determine protein expression of LDHC, integrin αv and integrin β3 in various cell lines. Approximately 5×10^5^ MDA-MB-468 cells were plated in 6-well plates, followed by incubation with CPP:siRNA (final siRNA conc = 200nM) for 72 hours after which protein lysates were isolated using RIPA buffer (Pierce, #89900) supplemented with a HALT protease and phosphatase inhibitor cocktail (Thermo Fisher Scientific, #87786). Protein lysates were centrifuged for 30 min at 20,000 x g and protein content of supernatants was determined using the BCA protein assay (Thermo Fisher Scientific, #23225). Protein samples were reduced and denatured in 4x Laemmli sample buffer (BioRad, #161-07470), loaded onto a 4–15% TGX gel (BioRad, #4561084) and transferred onto polyvinylidene fluoride (PVDF) membrane (BioRad, #1704156). Membranes were blocked in 5% non-fat dried milk/Tris-buffered saline with 0.1% Tween-20, washed and incubated overnight at 4 °C with the following primary antibodies diluted in blocking buffer; rabbit anti-human LDHC (Abcam, #ab52747, 1:1000), rabbit anti-human integrin αV (Cell signaling, #4711, 1:1000), rabbit anti-human integrin β3 (Cell signaling, #13166, 1:1000) and rabbit anti-human β-actin (Cell signaling, #4970, 1:1000). Next, membranes were washed, incubated with horseradish Peroxidase (HRP)-linked rabbit secondary antibody (Cell signaling, #7074, 1:5,000) for 1 h at room temperature, and proteins were detected by ECL Plus (Thermo Fisher Scientific, #32209) using the ChemiDoc XRS+ Imaging system (Bio-Rad). Images acquisition and densitometry analysis were performed using the IMAGE LAB software v6.1 (Bio-Rad).

### Cytotoxicity assay

To assess toxicity induced by CPP:siRNA treatment, breast cancer and non-cancerous cells were seeded at 1×10^4^ cells per well in an opaque 96-well plate and allowed to adhere overnight at 37 °C and % CO2. Next, 100µl CPP:siRNA complexes (50% v/v Opti-MEM I Reduced Serum medium and complete, antibiotic-free DMEM media) were added for 72 hours at 37°C and 5% CO2. Cytotoxicity was assessed using the CellTiter-Glo® reagent (Promega, #G7572) following the manufacturer’s guidelines and luminescence was recorded using the GloMax®-Multi Detection system (Promega).

### Clonogenic assay

To determine the effect of CPP:siRNA complexes on the long-term survival of breast cancer cells as single agents or in combination with Olaparib (Selleck Chemicals, #AZD2281), 1×10^4^ MDA-MB-468 breast cancer cells were seeded per well in a 12-well plate. On the 2^nd^ and 7^th^ day after seeding, cells were treated with CPP:siRNA complexes (50% v/v Opti-MEM I Reduced Serum medium and complete, antibiotic-free DMEM media) and on the 11^th^ day after seeding, cells were treated with olaparib (30 µM). The concentration of olaparib and treatment duration was chosen as previously described [10]. Then, on the 14^th^ day, cells were washed with DPBS (Gibco, #14190-094) and stained with 1% crystal violet (Sigma-Aldrich, #C6158) in 25% methanol. Excess stain was washed away, and crystal violet was eluted with 10% sodium dodecyl sulfate (SDS) and absorbance was measured at 590 nm on the NanoQuant infinite F200 Pro instrument (Tecan).

### Zebrafish maintenance and breeding

In vivo cytotoxicity and anti-tumor activity of CPP:siRNA complexes were determined using zebrafish. Wild type AB zebrafish (Danio rerio) were maintained in standard conditions at the zebrafish laboratory at Sidra Medicine, Doha, Qatar. Adult zebrafish were set up for breeding, embryos were collected and maintained in PTU-E3 media at 28.5 °C.

### Zebrafish embryo toxicity test

CPP:siRNA complexes (final siRNA concentration 150, 200, 250nM) were injected into single-cell stage wild type AB zebrafish embryos. At 4 days post-fertilization (dpf), the survival rate and morphology of the injected zebrafish embryos were compared to untreated embryos using the Zeiss Stemi 2000-C Stereo microscope (Zeiss). Zebrafish morphology was defined as G1: severely affected, G2: mildly affected, G3: normal.

### Breast cancer xenograft zebrafish model

The anti-tumor activity of the 10R-RGD:siRNA and cRGD-10R:siRNA complexes was assessed using a breast cancer xenograft zebrafish model. A total of 1 × 10^6^ MDA-MB-468 breast cancer cells were pre-labeled with Vybrant™ CM-DiI Cell-Labeling Solution (#V-22888, ThermoFisher scientific) for 20 min at 37°C. Next, 1×10^3^ Dil-labeled MDA-MB-468 cells (in 5nl complete DMEM media) were injected into the yolk sac of 48 hours post-fertilization (hpf) anesthetized zebrafish embryos using a Pico-Liter Microinjector. After microinjection, zebrafish embryos were maintained in PTU-E3 medium using 24 well plates at 34°C. After 24 hours post-injection (hpi), the embryos were imaged using the Zeiss AXIO Zoom.V16 microscope at 100x magnification and 560 nm, and embryos were selected based on size and location of tumor engraftment for further experiments. Next, CPP:siRNA complexes (final siRNA concentration 200nM in 25 µl of Ultra-Pure DNAse/RNAse free water with peptide:siRNA molar ratio of 5) were injected into the heart of the selected embryos at 32hpi. Finally, embryos were imaged at 72hpi using the Zeiss AXIO Zoom.V16 microscope at 100x magnification and 560 nm. A total of 60 z-stack images were acquired and processed into maximum intensity projection images using the ZEN black software. Average fluorescence intensity and area was determined using both ImageJ (v1.54g) and ZEN (v 3.10) software.

### Statistical analysis

Normality of data was assessed using the Shapiro–Wilk test, and the one-way analysis of variance (ANOVA) or two-tailed unpaired t-test were used to compare groups. P value ≤ 0.05 was defined as statistically significant. Data are represented as mean ± standard error of mean (SEM). Statistical analyses and data representation were performed using GraphPad prism v10.0.0 (San Diego, CA, USA).

## RESULTS

### Assessment of CPP:siRNA complex formation, serum stability and physicochemical characterization of the nanocomplexes

Gel retardation assays were used to determine the optimal ratio that is required to encapsulate the siRNA with minimum release and maximum binding ability. LDHC siRNA was incubated with each of the four CPPs at different peptide:siRNA molar ratios (2.5:1, 5:1, 10:1). As shown in **Figure 1A**, robust complex formation was observed for all CPPs at peptide:siRNA molar ratios of 5:1 and 10:1. The ability of the CPP:siRNA complexes to protect the siRNA from degradation was assessed using a serum stability assay (using 5:1 peptide:siRNA molar ratio), which showed that siRNA integrity was maintained over 24 hours in 50% human serum (**Figure 1B**).

**Figure 1.**
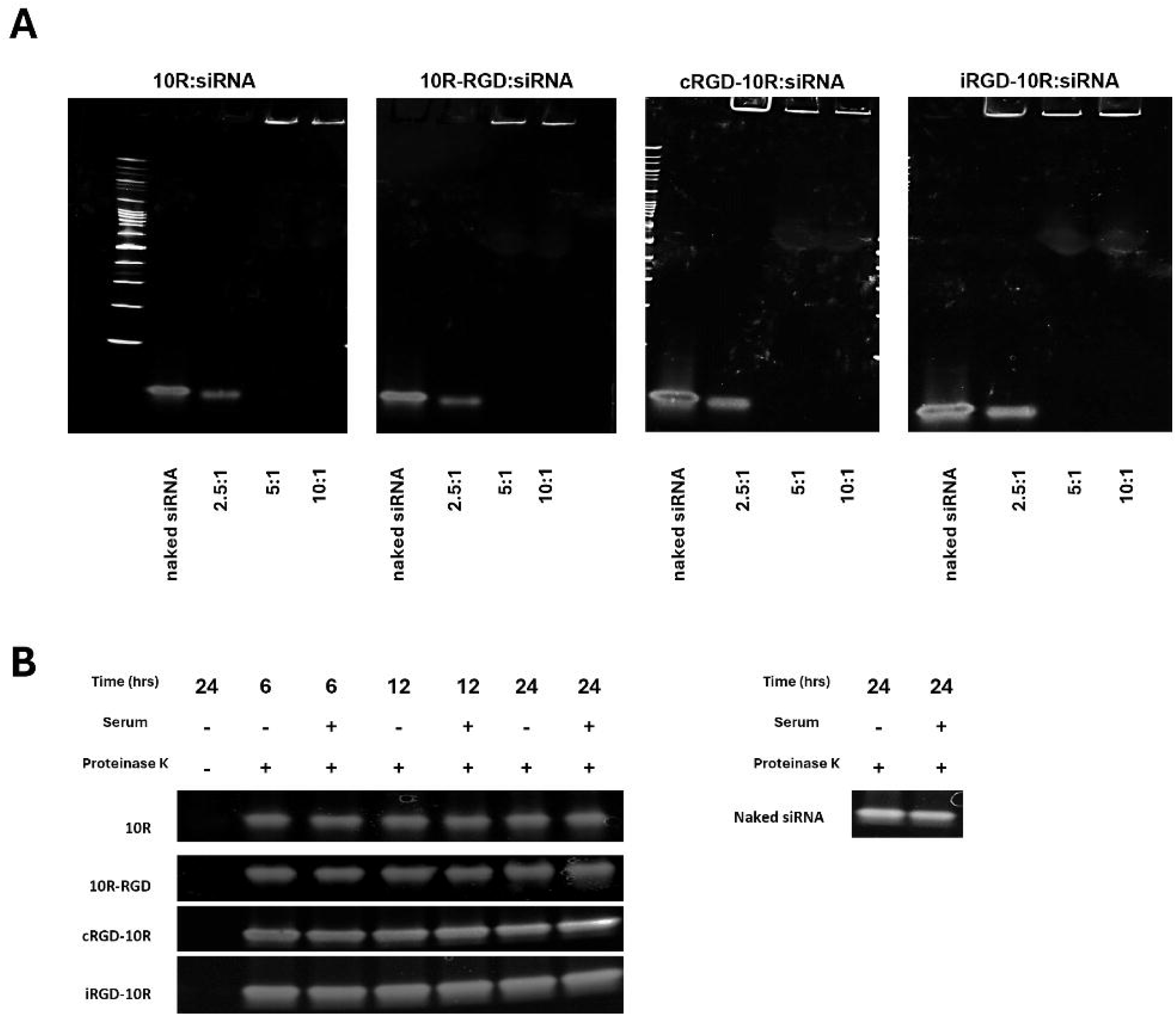
Assessment of CPP:siRNA complex formation and serum stability. (A) Visualization of 10R:siLDHC#2, 10R-RGD:siLDHC#2, cRGD-10R:siLDHC#2 and iRGD-10R:siLDHC#2 complex formation at different peptide:siRNA molar ratios using gel retardation assay. (B) Serum stability assay of CPP:siLDHC#2 complexes (5:1 peptide:siRNA molar ratio) in 50% human serum.

Dynamic light scattering (DLS) analysis showed that the four distinct CPP:siRNA complexes exhibit an average hydrodynamic diameter ranging between 129 and 168 nm, zeta potentials of 6.47±9.34 to 29.62±7.76 mV, and polydispersity indices below 0.25 (**Figure 2A**). Transmission electron microscopy (TEM) revealed the formation of uniform, circular nanocomplexes of expected, smaller size (56-95 nm) compared to hydrodynamic diameter measured by DLS (**Figure 2B**).

**Figure 2.**
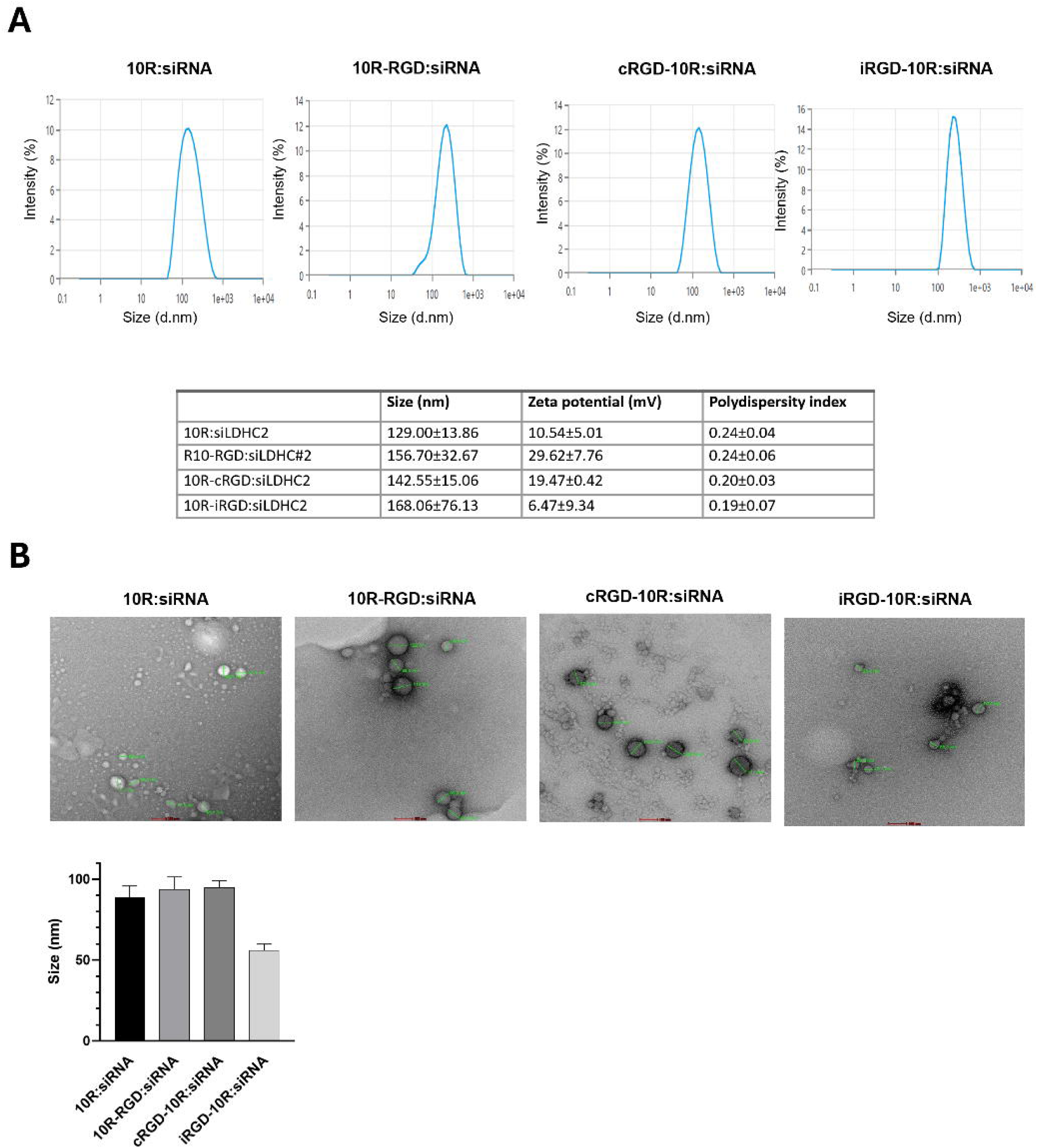
Physicochemical characterization of CPP:siRNA complexes. (A) Dynamic Light Scattering analysis of 10R:siLDHC#2, 10R-RGD:siLDHC#2, cRGD-10R:siLDHC#2 and iRGD-10R:siLDHC#2 complexes at peptide:siRNA molar ratio of 5:1. Complexes were diluted in Ultra-Pure DNAse/RNAse free water and analyzed at 677nm at room temperature with a constant angle of 90°. Values represent mean and standard error of mean (±SEM) from three independent replicates. (B) Representative negative stain TEM images of CPP:siLDHC#2 complexes. Bar chart represents means and standard error of mean (±SEM) from three independent experiments.

### CPP:siRNA complexes demonstrate good cellular uptake in breast cancer cells and exhibit favorable safety profile in vitro

We assessed the cellular uptake efficacy of the various CPP:siRNA complexes using integrin αvβ3 positive MDA-MB-468 breast cancer cells (**Figure 3**). The 10R-RGD:siRNA and cRGD-10R:siRNA complexes demonstrated the highest cellular uptake efficiency compared to the 10R:siRNA and iRGD-10R:siRNA complexes. Next, a range of triple negative breast cancer cell lines and two non-cancerous cell lines were used to investigate the potential presence of toxicity following treatment with the CPP:siRNA complexes. The cell lines were chosen based on their expression of integrins αv and β3 **(Figure S1)** to allow the simultaneous assessment of toxicity related to unspecific uptake or RGD-mediated cellular uptake. The breast cancer cell lines MDA-MB-468 and BT-549 were selected as integrin αvβ3 positive cancer cell line models, while the MDA-MB-453 and DU4475 breast cancer cell lines were chosen to represent integrin αvβ3 negative cancer cells. In addition, the integrin αvβ3 expressing IMR-90 and HUVEC cells were used to study the safety profiles of the CPP:siRNA complexes in non-cancerous cells. No significant cytotoxicity was observed in either the breast cancer cell lines or the non-cancerous cells, except for a minor toxicity of the 10R-RGD:siRNA complex in MDA-MB-468 cells (12% toxicity) and DU4475 cells (20% toxicity), suggesting that our CPP:siRNA complexes overall exhibit a favorable safety profile (**Figure 4**). Lipofectamine-mediated cellular uptake of naked siRNA induced toxicity to varying degrees as commonly seen using liposome transfection [20].

**Figure 3.**
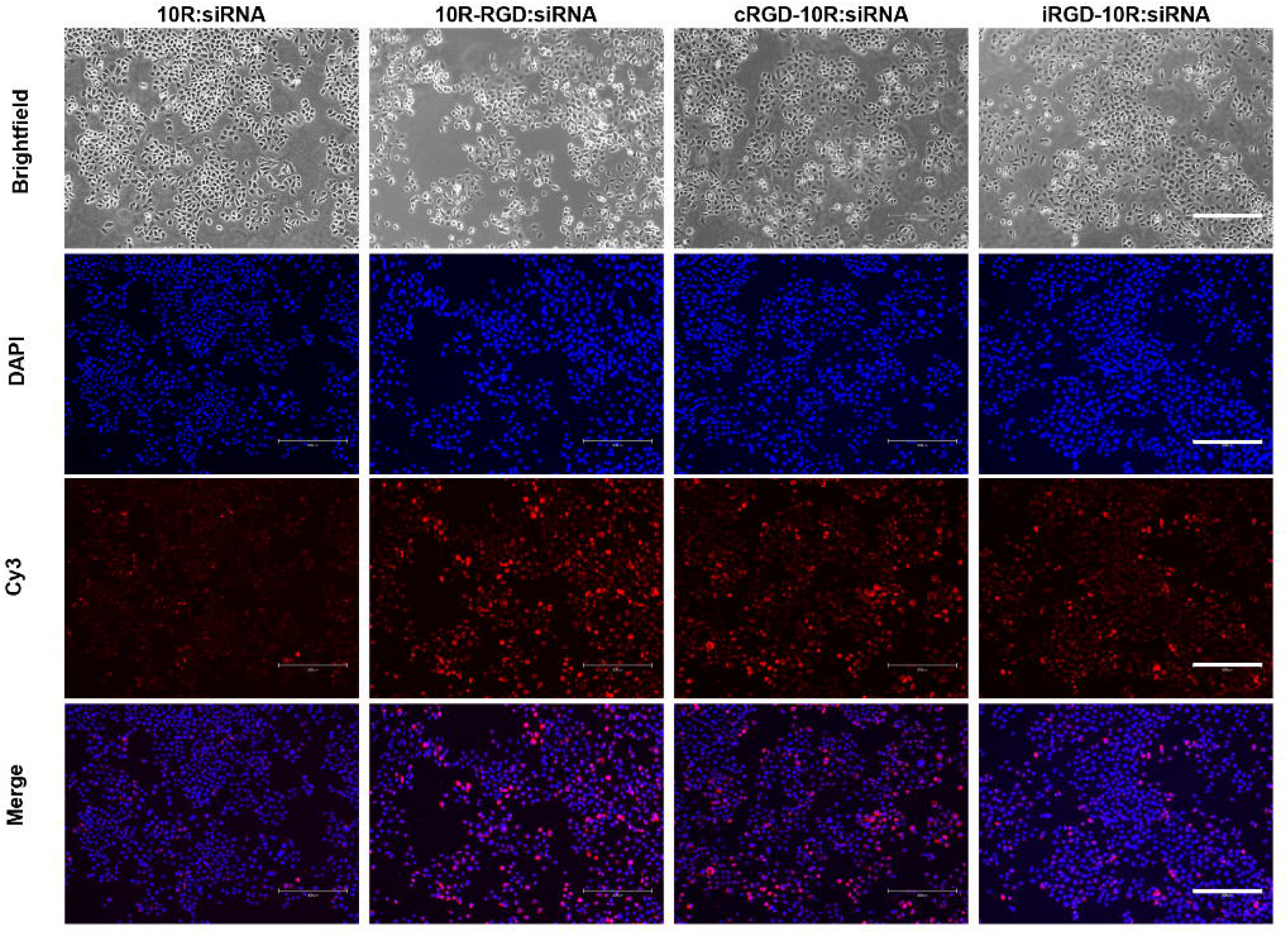
Cellular uptake of CPP:siRNA complexes in MDA-MB-468 breast cancer cells. Immunofluorescent imaging of Cy3-prelabeled CPP:siRNA complexes after 72 hours incubation. Blue, DAPI; red, Cy3-labeled siRNA.

**Figure 4.**
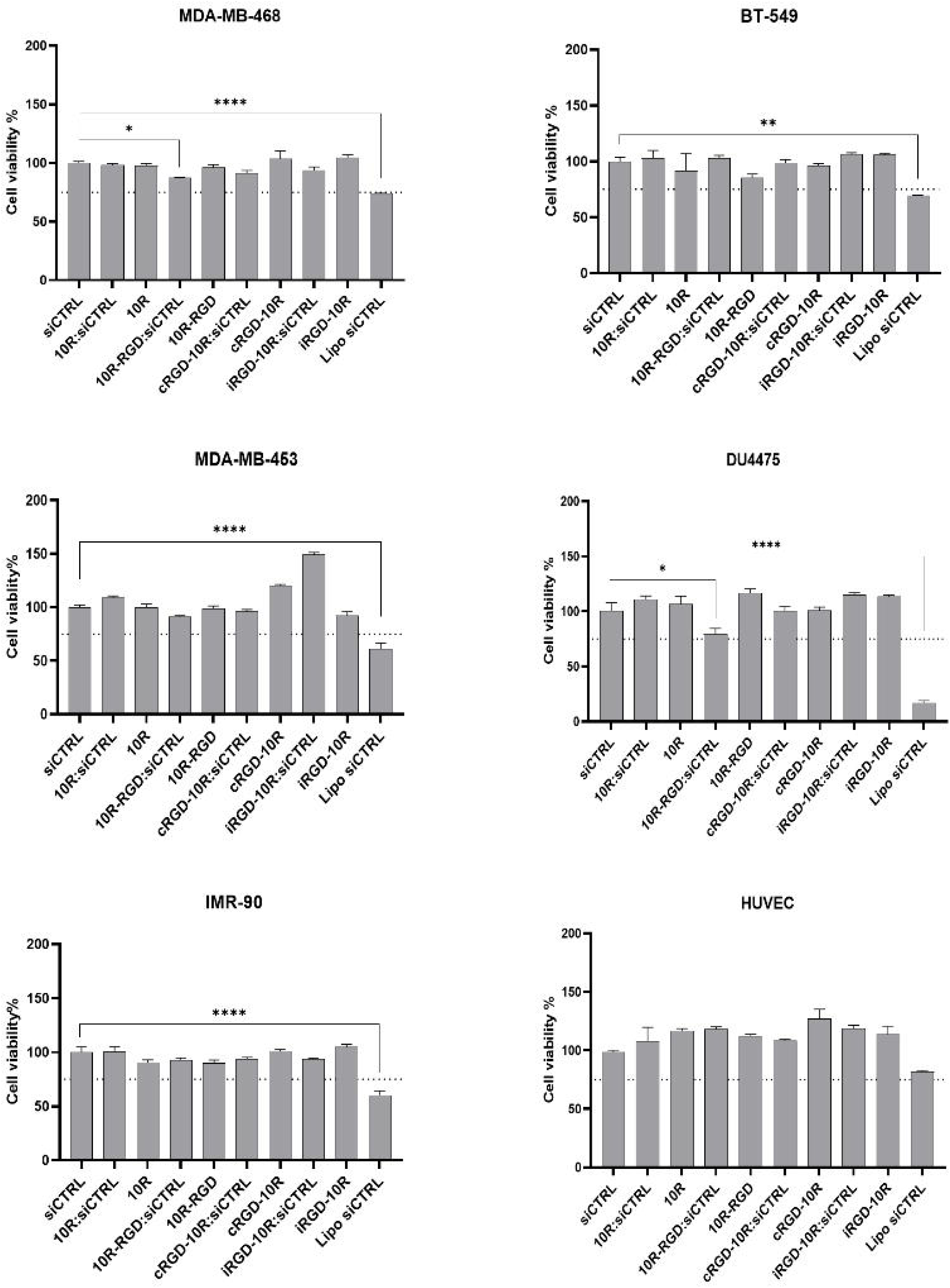
Treatment with CPP:siRNA complexes is associated with a favorable safety profile in vitro. Breast cancer and non-cancerous cell lines were treated with CPP:siRNA complexes (5:1 molar ratio) for 72 hours, and cytotoxicity was determined using CellTiter-Glo luminescent cell viability assay. Lipofectamine-mediated transfection of naked siCTRL was used as a positive control for efficient cellular uptake, and CPPs alone were used to assess the toxicity of the peptides. Error bars represent standard error of mean (±SEM) from three independent experiments. Statistical analysis to assess reduction in cell viability was performed using the one-way ANOVA test with Dunnett correction. * p<0.05, ** p<0.01; **** p<0.0001.

These findings are in line with previous studies reporting low cytotoxicity levels using CPPs [21,22].

### siRNA delivery by CPPs efficiently reduces LDHC expression and clonogenic ability in triple negative breast cancer cells in vitro

Given the robust complex formation, good cellular uptake and low cytotoxicity of the CPP:siRNA complexes, we next sought to evaluate their LDHC knockdown efficiency in the integrin αvβ3 positive, LDHC expressing triple negative breast cancer cell lines MDA-MB-468 and BT-549.We found that the 10R, 10R-RGD and cRGD-10R peptides complexed with siLDHC#2 significantly reduced *LDHC* mRNA expression in MDA-MB-468 cells, with a borderline reduction in *LDHC* expression using the iRGD-10R peptide (**Figure 5A**). On the other hand, more modest reductions in *LDHC* expression were seen in BT-549 cells (**Figure 5B**), which is likely the result of their lower endogenous LDHC expression and likely reduced silencing efficiency [10]. Notably, we obtained a greater reduction in *LDHC* expression using the CPP:siRNA complexes as compared to the naked siRNA. Furthermore, the knockdown efficiency of the CPP:siLDHC complexes was confirmed at protein level with the 10R-RGD:siLDHC#2 and cRGD-10R:siLDHC#2 complexes showing the highest efficiencies, reducing LDHC protein expression by 38% and 41% respectively (**Figure 5C**). The clonogenic assay was used to evaluate the effect of CPP-mediated LDHC silencing, LDHC in combination with olaparib, on the long-term survival of MDA-MB-468 cells. Based on the knockdown efficiency results, the 10R-RGD:siRNA and cRGD-10R:siRNA complexes were selected for further functional validation. In accordance with our previous observations [10], CPP-mediated silencing of LDHC alone significantly reduced the clonogenic ability of MDA-MB-468 triple negative breast cancer cells and improved treatment response to olaparib (**Figure 6**). This decrease in clonogenicity highlights the therapeutic potential of targeting LDHC in TNBC.

**Figure 5.**
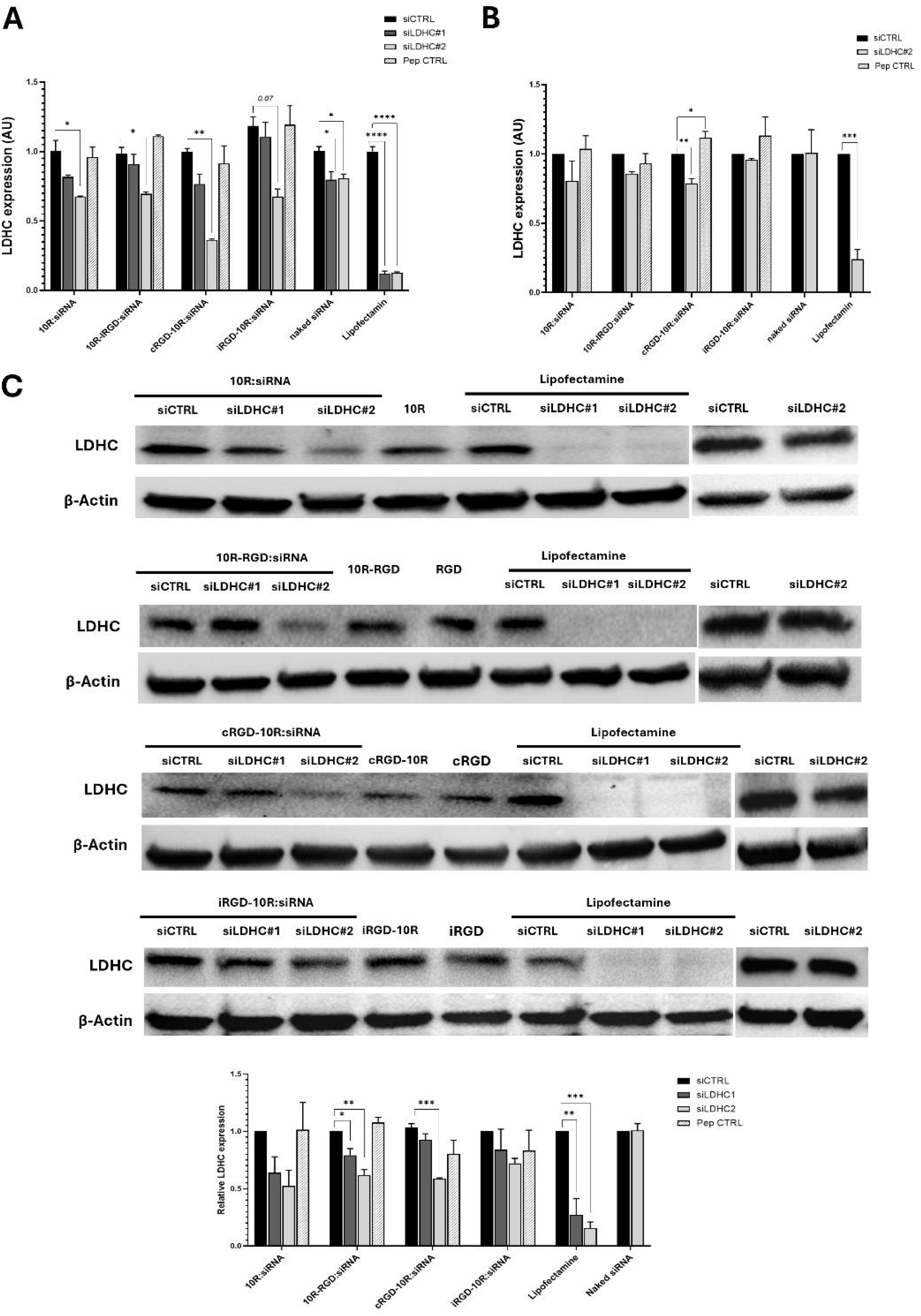
CPP:siRNA complexes demonstrate efficient LDHC knockdown in triple negative breast cancer cells in vitro. (A) LDHC mRNA expression, normalized to RPLPO, of MDA-MB-468 and (B) BT-549 cells following treatment with CPP:siRNA complexes for 72 hours. (C) Representative images of LDHC protein expression of MDA-MB-468 cells treated with CPP:siRNA complexes for 72 hours. β-actin was used as loading control. Bar chart represent densitometry values from three independent experiments (mean±SEM). Statistical analysis was performed using the one-way ANOVA test with Dunnett correction for comparison of more than two groups and two-tailed unpaired t-test for comparison of two groups. * p<0.05, ** p<0.01, *** p<0.001, **** p<0.0001.

**Figure 6.**
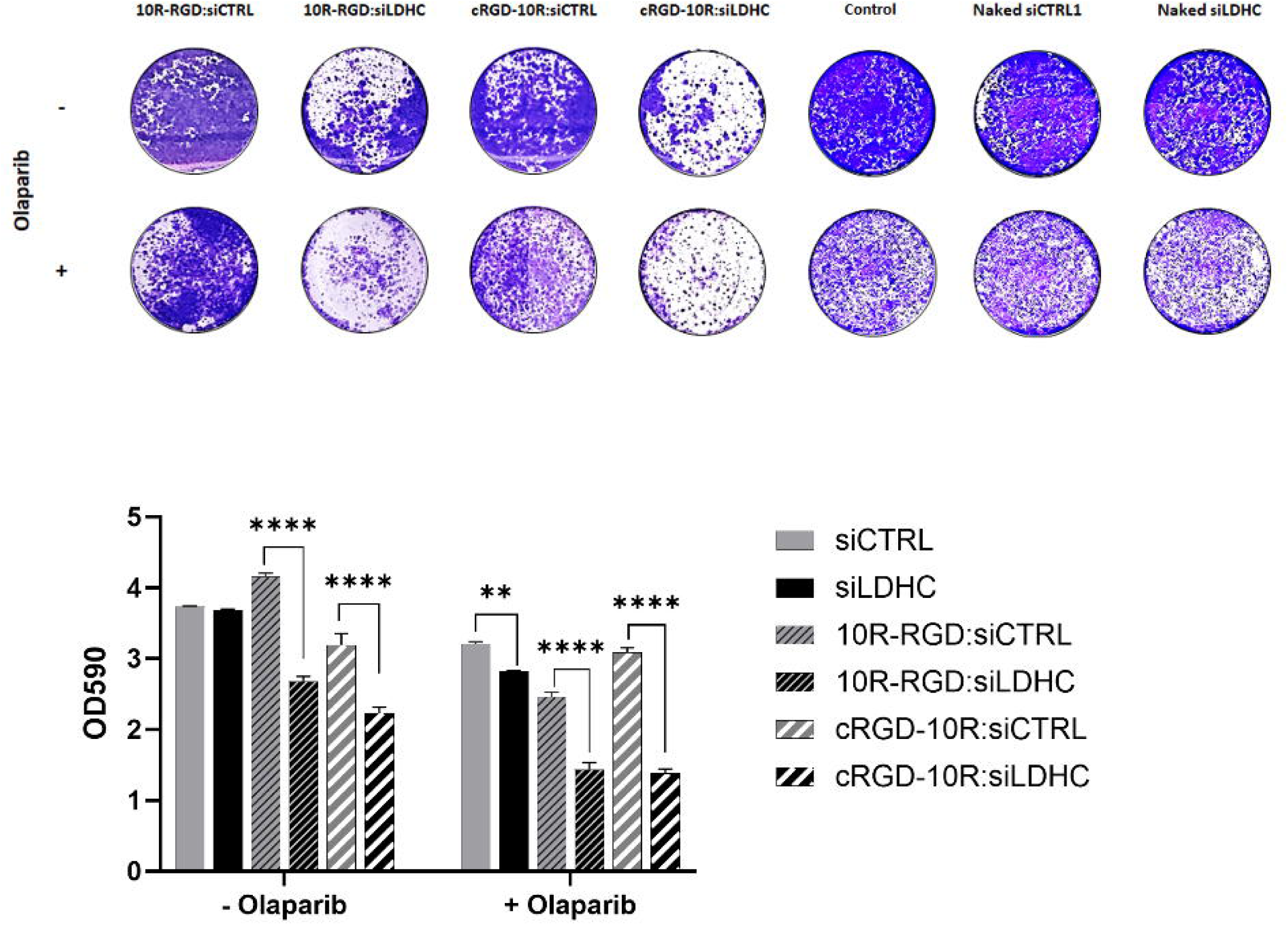
CPP:siLDHC treatment significantly reduces the clonogenic ability of MDA-MB-468 cells and potentiates olaparib treatment. Representative images of clonogenic assay using 10R-RGD:siRNA and cRGD-10R:siRNA alone or in combination with olaparib short-term treatment (72 hours). Bar chart depicting crystal violet absorbance measurements from three independent experiments (mean ±SEM). Statistical analysis was performed using the one-way ANOVA test with Šídák correction. * p<0.05, ** p<0.01, *** p<0.001.

### 10R-RGD and cRGD-10R:siRNA complexes exhibit anti-tumor activity with minor toxicity in breast cancer xenograft zebrafish model

Our promising in vitro results, showing good LDHC knockdown efficiency with reduced colony-forming ability and low cytotoxicity, prompted us to evaluate the toxicity of the CPP:siRNA nanocomplexes *in vivo*. Cytotoxicity was assessed through microinjection of 10R-RGD:siRNA and cRGD-10R:siRNA complexes (150, 200 and 250nM) into single-cell stage wild type AB zebrafish embryos. Minor toxicity was observed for both CPP:siRNA complexes at various doses (**Figure 7A**). No significant morphological abnormalities of the zebrafish embryos were observed after microinjection with either 10R-RGD:siRNA or cRGD-10R:siRNA complexes (**Figure 7B**).

**Figure 7.**
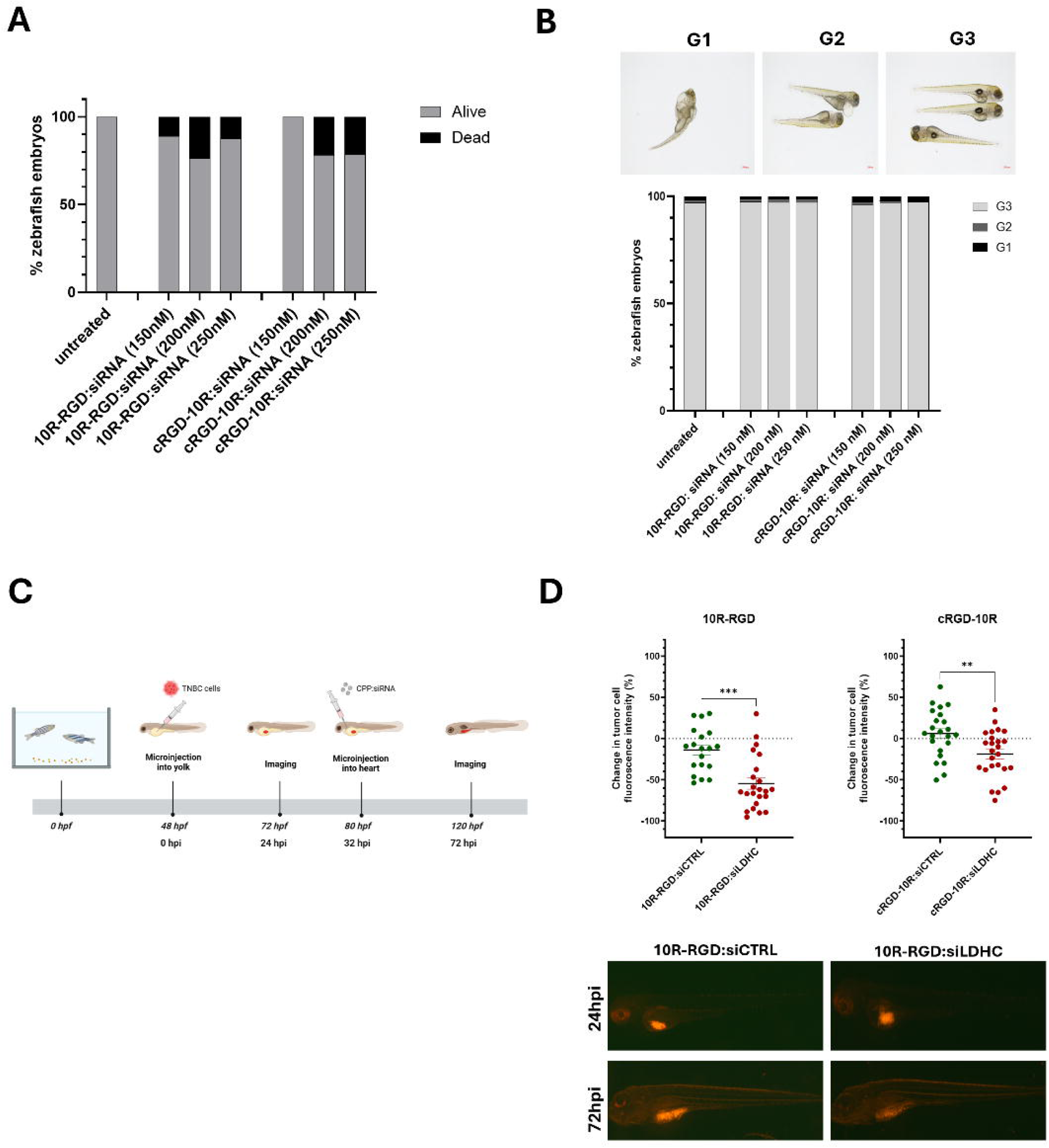
10R-RGD and cRGD-10R:siRNA complexes exhibit anti-tumor effects in TNBC xenograft zebrafish model without inducing toxicity. (A) Zebrafish mortality and (B) morphological abnormalities following treatment with 10R-RGD:siRNA and cRGD-10R:siRNA complexes. G1: severely affected, G2: mildly affected, G3: normal. (C) Diagram of TNBC xenograft zebrafish model, depicting TNBC cell injection in the yolk at 2 days post-fertilization (dpf) and CPP:siRNA intracardial treatment at 32 hours post-injection (hpi). (D) Change in tumor cell burden, measured as % change in fluorescence intensity following CPP:siRNA treatment. Representative images of fluorescent tumor cells at 24hpi and 72hpi. Scatter dot plots represent fluorescence intensity values from two independent experiments (mean ±SEM). Statistical analysis was performed using the two-tailed unpaired t-test. ** p<0.01, *** p<0.001.

Next, the therapeutic potential of CPP-based siLDHC delivery was assessed using a TNBC xenograft zebrafish model (**Figure 7C**). Treatment of MDA-MB-468 xenograft zebrafish larvae with cRGD-10R and 10R-RGD:siLDHC nanocomplexes resulted in a significant reduction in tumor burden (**Figure 7D**), achieving up to 50% reduction in tumor size using the 10R-RGD complex. These findings are in line with our in vitro results that demonstrate a higher cellular uptake and *LDHC* knockdown efficiency using 10R-RGD:siRNA complexes compared to cRGD-10R:siRNA.

## DISCUSSION

Over the last few decades, cancer treatment has made great strides with targeted therapy and immunotherapy improving the prognosis of patients with various cancer types. Targeted therapy with mTOR, CDK4/6 or PI3K inhibitors in combination with hormonal therapy has now entered the clinic for the treatment of advanced breast cancer patients with hormone receptor positive tumors [23]. Further, patients with Her2-enriched breast tumors benefit from combination treatment of chemotherapy with a variety of anti-Her2 treatment modalities [24]. Despite these advances in molecular targeting, treatment options for patients with triple negative breast cancer remain limited. While TNBC patients with *BRCA1/2* mutations may benefit from PARP inhibitors, and the prognosis of patients with PD-L1 tumor expression may be improved by immune checkpoint blockade, the majority of TNBC patients still receive standard-of-care chemotherapy alongside radiotherapy and surgery [2,25]. Moreover, while TNBC patients often exhibit good pathological response rates following chemotherapy, their overall prognosis remains poor due to early cancer recurrence and the development of chemoresistance. Hence, there is an unmet need to find novel therapeutic targets to increase clinical outcomes for these patients.

Previously, we found that targeting LDHC, a highly tumor-specific metabolic enzyme, significantly augments genomic integrity, reduces the clonogenic ability of breast tumor cells and enhances treatment responses to DNA damage repair-related drugs *in vitro*. Hence, we hypothesized that targeting of LDHC could be used to complement traditional cancer treatments to enhance anti-tumor efficacy. However, specific LDHC inhibitors and blocking antibodies are currently unavailable. Therefore, we explored the therapeutic potential of siRNA-based drugs to target LDHC tumor expression. To date, five siRNA-based drugs have been approved by the FDA for the treatment of liver diseases, of whom four utilize N-acetylgalactosamine (GalNAc) as a targeting ligand for the asialoglycoprotein receptor (ASGPR), predominantly expressed in hepatocytes [26]. Although no siRNA therapeutics have been approved for cancer treatment to date, numerous candidates are currently in phase I/II clinical trials [26].

In the present study, we used cell penetrating peptides as a delivery system for LDHC siRNA in triple negative breast cancer cells. More specifically, we explored the use of the integrin αVβ3 RGD binding motif to facilitate tumor homing and penetrance of the siRNA therapeutic. Four CPPs; 10R, 10R-RGD, cRGD-10R and iRGD-10R; were studied for siRNA complexing efficiency, serum stability and cytotoxicity *in vitro*. All four CPPs formed uniform structures with the siRNA molecules, resulting in positively charged nanocomplexes which enhance permeability and retention [27]. Complexing siRNA with the CPPs enhanced their persistence in human serum, indicating that the peptides were able to protect the siRNA from circulating RNA enzymes. Furthermore, no to minor toxicity was observed using the CPP:siRNA complexes in either breast cancer cells (αVβ3 negative or positive TNBCs) or non-cancerous cells (integrin αVβ3 positive IMR-90 and HUVEC cells), indicating a favorable safety profile with no adverse side effects. Comparative analysis of LDHC knockdown efficiency revealed that the 10R-RGD:siLDHC and cRGD-10R:siLDHC complexes more effectively reduced LDHC expression in MDA-MB-468 triple negative breast cancer cells (integrin αVβ3 positive), achieving up to a 40% reduction at the protein level and 63% at the mRNA level. These findings are in accordance with other preclinical studies reporting silencing efficiencies of approximately 70% using CPPs in cancer. For example, Van Asbeck et al reported knockdown efficiencies ranging from 70-85% using PF6 and PF14 encapsulated siRNA, while C6M1 conjugated siRNA targeted GAPDH in ovarian cells with a knockdown efficiency of about 70% [28,29].

Next, we evaluated the therapeutic potential of targeting LDHC in TNBC using CPP-siRNA based therapy. Based on our previous work, we investigated whether CPP:siLDHC treatment alone and in combination with olaparib impacts the clonogenic ability of TNBC cells. Similarly to our previous observations, CPP:siRNA-based targeting of LDHC, in particular using 10R-RGD:siLDHC and cRGD-10R:siLDHC, significantly reduced tumor cell clonogenic ability and greatly enhanced the anti-tumor effect of olaparib, indicating that CPP:siLDHC therapy could provide a promising strategy for the treatment of TNBC. To validate the clinical significance of CPP:siLDHC-based therapy, the anti-tumor activity and safety profiles of 10R-RGD:siLDHC and cRGD-10R:siLDHC delivery were investigated in a TNBC xenograft zebrafish model. Both complexes significantly reduced tumor burden, with the 10R-RGD complex achieving up to a 50% reduction in tumor size, without inducing major morphological abnormalities or death.

## CONCLUSION

In conclusion, we identified two CPP:siRNA nanocomplexes; 10R-RGD:siLDHC and cRGD-10R:siLDHC; that persist in serum and effectively target LDHC in triple negative breast cancer, exhibiting significant anti-tumor activity and a favorable safety profile in vitro and in vivo. These findings corroborate LDHC as a promising target for TNBC therapy and highlight the potential of CPP:siRNA-based drugs as a novel therapeutic approach for cancer therapy.

## DECLARATIONS

### ETHICS APPROVAL AND CONSENT TO PARTICIPATE

All animal work was performed according to the Ministry of Public Health (MOPH), Qatar animal research guidelines under an approved protocol by the Institutional Animal Care and Use Committee (protocol SIDRA-2024-005), and adhered to the ARRIVE guidelines.

### AVAILABILITY OF DATA AND MATERIALS

All data that supports the findings of this study are included in this published article. Additional data can be provided upon request from the corresponding author.

## ACKNOWLEDGEMENTS

We would like to thank Dr Sahar Da’as and her team from the Zebrafish Core Facility, Sidra Medicine, Qatar for their invaluable contributions to the xenograft zebrafish work. We would like to acknowledge the HBKU Materials Core Lab for their valuable support and resources provided.

## AUTHOR CONTRIBUTIONS

HQ: Formal Analysis, Investigation, Methodology, Visualization, Writing – original draft, Writing – review & editing; AN: Formal Analysis, Investigation, Methodology, Visualization, Writing – review & editing; TG: Formal Analysis, Writing – review & editing; JP: Formal Analysis, Writing – review & editing; UJ: Formal Analysis, Investigation, Writing – review & editing; MS: Investigation, Writing – review & editing; RT: Investigation, Writing – review & editing; KAM: Methodology, Writing – review & editing; JD: Conceptualization, Formal Analysis, Funding acquisition, Project Administration, Supervision, Visualization, Writing – original draft, Writing – review & editing. All authors read and approved the final manuscript.

