## Supplementary figures and images for "Anti-tumor effects of a novel cell penetrating peptide-based therapeutic approach to target Lactate Dehydrogenase C (LDHC) in triple negative breast cancer"

### Figure S1

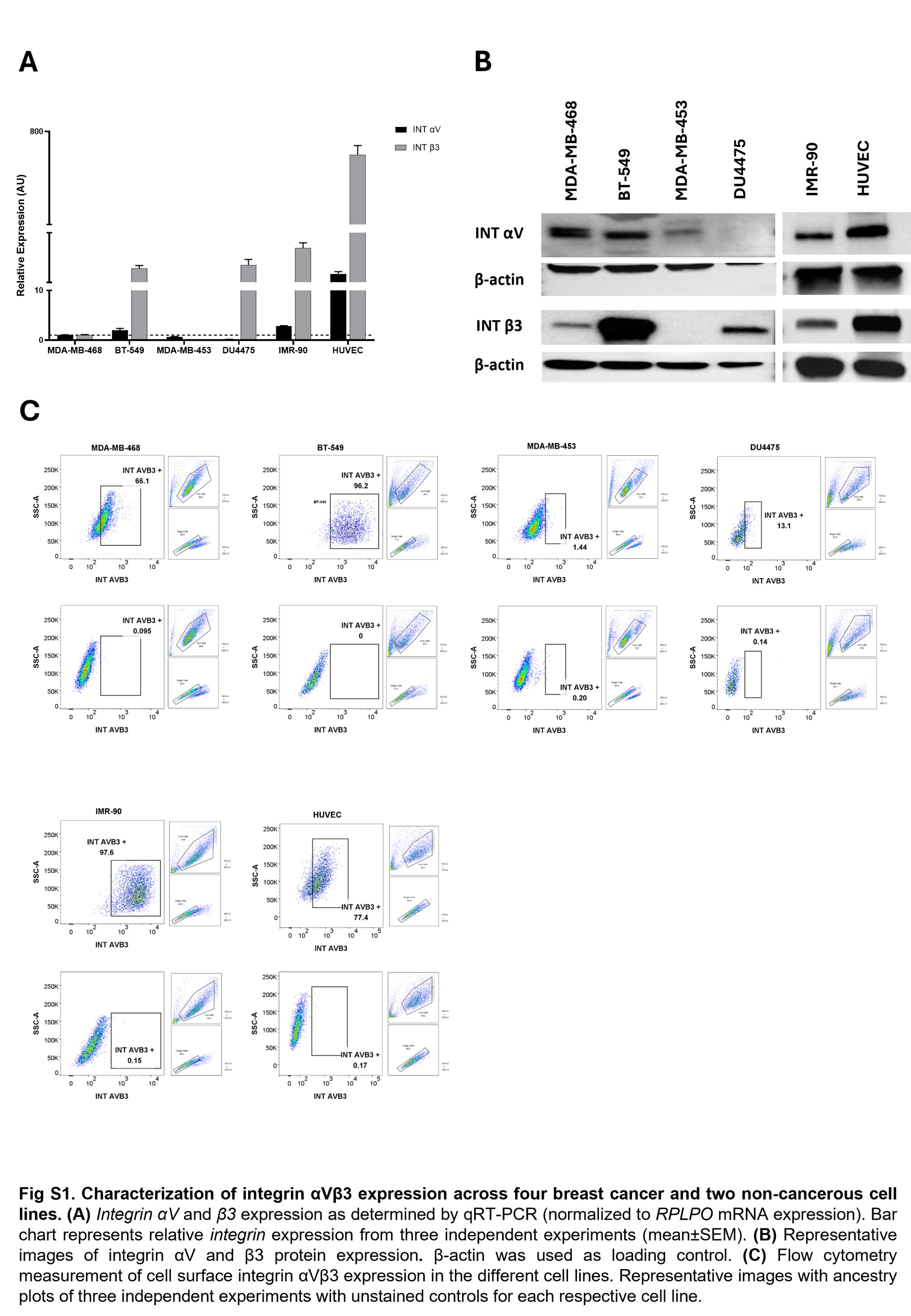
